# Deep Phenotyping with Global Brain Activity and Plasticity Mapping Identify the Dorsal Raphe-Basolateral Amygdala Circuit as a Mediator of Adaptive Stress Responses

**DOI:** 10.64898/2026.07.24.740522

**Authors:** Shiladitya Mitra, Tibor Stark, Marcin Baranski, Beatrice Dal Bianco, Carlo Castoldi, Maddalena Pieroni, Sowmya Narayan, Cristina Beer, Rosa Eva Huettl, Monika Pawlowska, Lotte van Doeselaar, Joeri Bordes, Margherita Springer, Huanqing Yang, Veronika Kovarova, London Aman, Benjamin Jurek, Akshaya Rajan, Nicolas Snaidero, Michael Czisch, Marzena Stefaniuk, Bianca A. Silva, Mathias V. Schmidt

**Author notes:** Correspondence: Mathias V. Schmidt, Research Group Neurobiology of Stress Max Planck Institute of Psychiatry Kraepelinstr. 2-10, 80804 Munich Germany. Co-correspondence: Shiladitya Mitra.

## Abstract

Exposure to chronic environmental challenges triggers divergent behavioral trajectories across individuals. At the core, these different trajectories can be classified as individuals actively adapting to the challenges (“responders”) and those displaying a rigid, non-responsive phenotype (“non-responders”). The brain system-wide network configurations that dictate why individuals diverge along these differential coping strategies, which can also lead to disease vulnerability or resilience, remain poorly understood. Here, we paired machine-learning-based deep behavioral phenotyping with multi-modal whole-brain imaging, integrating longitudinal Manganese-Enhanced MRI (MEMRI) and post-challenge cFOS mapping, to chart the functional landscape of individual stress trajectories in mice subjected to chronic social defeat stress. High-dimensional behavioral phenotyping revealed that active stress adaptation is a complex trajectory marked by latent, pre-stress kinetic signatures in vigilance-like and locomotive behaviors. At the neural level, longitudinal MEMRI captured distinct, consolidated activity reconfigurations across canonical valence and stress-regulatory circuits that segregated responders from non-responders. Complementary whole-brain cellular cFOS network analysis after an additional acute challenge revealed that non-responders exhibited marked hyper-modularity and network fragmentation, whereas responders feature a tightly integrated functional module co-clustering the periaqueductal gray, ventral tegmental area, basolateral amygdala (BLA), and dorsal raphe (DR). Notably, functional network connectivity along the DR-BLA axis was completely lost in non-responsive animals. Finally, pathway-specific chemogenetic inhibition of BLA-projecting DR neurons during a social challenge significantly attenuated social avoidance and reversed anxiety-like behavioral deficits, effectively shifting active behavioral adaptation toward a non-responsive phenotype. Together, these findings demonstrate that individual stress-coping strategies are driven by coordinated, system-wide reconfigurations of activity and plasticity, identifying the DR-BLA circuit as a critical gatekeeper of adaptive stress responses.

**Graphical Abstract:** Global neural functional alterations defining responding vs non-responding populations following chronic stress are understudied, yet crucial. Deep phenotyping followed by mapping brain-wide activity and plasticity changes identified these underlying divergent functional networks. Acute manipulation of a dorsal raphe – basolateral amygdala pathway ameliorated adaptive stress responses, highlighting the significance of this network-based approach.

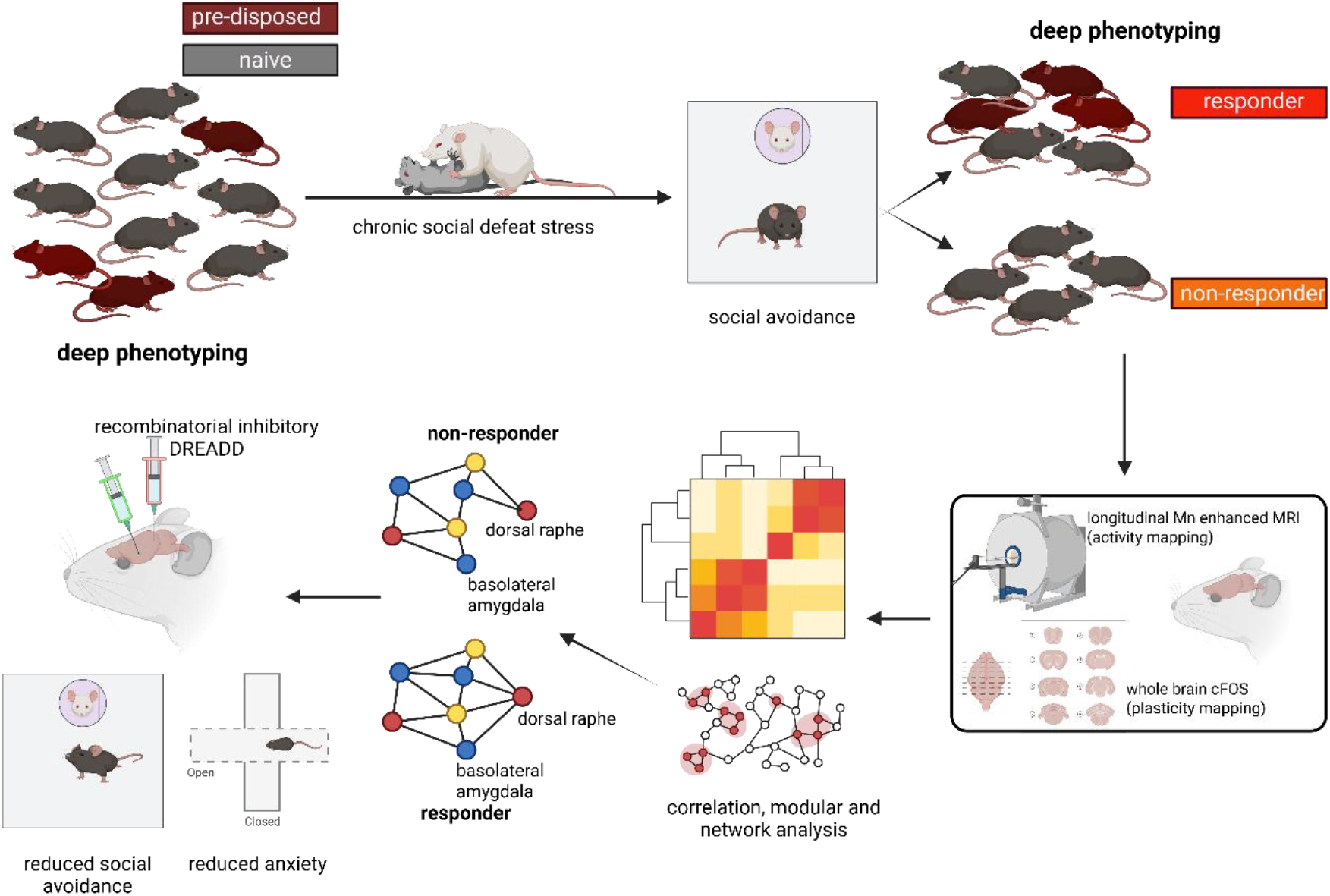

## Introduction

Exposure to environmental challenges triggers a suite of responses and physiological adaptations designed to maintain homeostasis and successful stress coping. However, the trajectory of these responses varies significantly across individuals. While some maintain functional equilibrium, others develop maladaptive phenotypes that mirror human neuropsychiatric conditions (1,2). This spectrum of individual variability is well-documented in both clinical populations and rodent models (3–7). While decades of research have utilized standardized stress paradigms to map the molecular and functional changes underlying these responses, a multimodal approach for a consolidated holistic portrayal of the changes in the systemic, brain-wide networks that dictate why individuals diverge in their stress responses remains incompletely represented.

The divergence in stress responses typically result in two distinct phenotypic strategies. In some individuals, exposure to stressors triggers an active adaptation of physiology and behavior to the challenging environment. Conversely, others display a more rigid phenotype, remaining largely unaffected by the changing environmental demands (1,7,8). Though these divergent trajectories are shaped by a complex interplay of genetic predisposition and prior experience, significant individual variability emerges even within standardized preclinical cohorts. This suggests that factors such as individual perception of the stressor and underlying epigenetic modifications also play decisive roles in shaping the eventual phenotype (5,9,10). Understanding how these distinct strategies are encoded requires a shift from examining isolated molecular changes to identifying the macroscopic patterns of neural activity that govern why one individual adapts while another remains resistant to change (11).

Examination of the neural signatures underlying differential stress coping strategies have traditionally been studied with a heavy bias toward a few canonical regions, such as the prefrontal cortex, nucleus accumbens and hippocampus (1,12,13). However, given the systemic nature of the stress response, focusing on isolated structures or circuits may overlook the broader functional networks that coordinate these phenotypes. There is a growing need for unbiased, whole-brain investigations that can capture the global landscape of neural activity and plasticity in relation to divergent stress responses (7,14–18). Furthermore, identifying these patterns requires a detailed behavioral phenotyping approach that provides nuanced differences beyond traditional behavioral metrics. By utilizing high-resolution, automated behavioral tracking, it becomes possible to capture the subtle, nuanced kinetics of social and non-social behaviors that distinguish an actively adapting individual from one displaying a rigid, non-responsive phenotype (19,20).

To address these questions, we utilized the Chronic Social Defeat Stress (CSDS) paradigm to induce divergent behavioral trajectories in mice (21). In addition to relying on traditional binary metrics (22), we employed a deep-learning-based framework—integrating DeepLabCut and DeepOF—to perform high-dimensional phenotyping of social interactions (23,24). This allowed us to precisely characterize the nuanced behavioral kinetics of individuals that actively adapt to stress (responders) compared to those displaying a rigid phenotype (non-responders). To map the neural correlates of these states, we implemented a multi-modal imaging approach: Manganese-Enhanced MRI (MEMRI) was used to capture longitudinal changes in *in vivo* brain activity, complemented by *post hoc* whole-brain cFOS immunolabeling to assess global plasticity. This unbiased screen identified, among multiple network alterations, a conserved functional network centred on the dorsal raphe (DR) to basolateral amygdala (BLA) circuit. Finally, using chemogenetic inhibition, we demonstrated that modulating this DR-BLA pathway was sufficient to attenuate the adaptive stress response and a responsive phenotype. Together, our findings provide a comprehensive, brain-wide map of activity and plasticity changes that accounts for stress-induced variability and identifies a critical circuit-level target for therapeutic intervention in stress-vulnerable populations.

## Material and Methods

### Animal Handling and breeding

Adult C57Bl/6n inbred mice (postnatal day (PND 60-90), mature adult CD1 mice (PND>180) and juvenile CD1 mice (PND 30-42) were used in the experiments. Only male mice have been used for the social defeat stress paradigm. These mice were obtained from Charles River and were bred and maintained in-house under standard conditions in individual ventilated cages (12h/12h light/dark cycle, 55% humidity and 23 ± 2°C). Food (regular chow diet) and water were supplied ad libitum. Mice were group-housed unless required by experimental procedures. Mice were acclimatized to the animal facility for at least a week before the start of the experiments.

All protocols were approved by the committee of the Care and use of Laboratory Animals of the government of Upper Bavaria, Germany and all procedures were in accordance with the European Coummunities Council directive 2010/63/UE of the European Parliament and of the Council of 22 September 2010 on the protection of animals used for scientific purposes.

### Chronic Social Defeat Stress (CSDS) paradigm

A 10 days chronic social defeat stress (CSDS) paradigm was employed to induce stress in mice and was similar to those described before (22,23). Briefly, 2-3 months old male C57Bl/6n mice were introduced in a cage containing an aggressive mature male CD1 mouse for 10 minutes every day for ten days. At the end of daily defeat, the C57Bl/6n mice were housed in the same cage next to the aggressor CD1 separated by a transparent barrier with perforations to enable visual and olfactory contact. The CD1 mice had been selected based on aggressive behavior and care was taken so that no severe wound was inflicted during the defeat procedure. In case of a fight, it would be broken by tapping on the grill of the cage. In instances where the fight is violent, mice were separated physically. A Latin square approach was used to prevent habituation and to randomize the aggressor every day. Different cohorts were used for experiments having different objectives – deep phenotyping analysis, MEMRI and global cFOS analysis.

### Behavior Tests

#### Automated behavior analysis

Behavioral Tests were performed in the morning after the CSDS paradigm on the 11^th^ and the 12^th^ day for social avoidance (SA) and elevated plus maze (EPM) test, respectively. These behavioral tests were recorded and analysed using Anymaze (Stoetling).

#### Social Avoidance Test

Social Avoidance (SA) test was conducted based on a previously described procedure (22,23). Briefly, C57Bl/6n mouse was placed in a square arena 40×40cms containing a small enclosure near one of the walls with or without a male mature CD1 mouse restricted in it. The C57Bl/6n mouse was allowed to explore for 5 minutes, after which a male mature CD1 mouse was introduced in the enclosure and the test continued for another 5 minutes. Time spent in a virtual ‘interaction zone’ near the enclosure and away from it was measured. Social avoidance interaction ratio was calculated based on time spent in the interaction zone with the CD1 mouse or without. C57Bl/6n mice subjected to the CSDS stress paradigm were categorized into non-responder or responders based on a z-score derived from time spent near the enclosure containing the aggressor CD1 mouse and the social interaction ratio.

#### Elevated Plus Maze Test

Elevated Plus Maze (EPM) test was conducted following the procedure as described earlier (25,26). Briefly C57Bl/6n mice were placed in the centre of the EPM arena facing the open arm and allowed to explore for 5 minutes. The time spent in each of the arms and the frequency of visits were analysed.

#### Deep learning based behavioral analysis

##### Free Social Interaction Test

Mice were subjected to a social interaction test before and after the CSDS paradigm post traditional behavioral tests SA and EPM. C57Bl/6n mouse was introduced into a cylindrical arena of 50cm diameter and was allowed to habituate for 10 minutes following which a male juvenile CD1 mouse was introduced. Interaction over the course of 10 minutes was recorded for subsequent deep phenotyping analysis.

##### Deep Phenotyping analysis

We employed automated pose estimation software DeepLabCut v2.2.2 for video analysis and utilized deep learning-based motion tracking and social interaction analysis tool DeepOF v07.0.2 for deep phenotyping as described earlier (23,27). We categorized behavioral outcomes into principal component analysis (PCA) to reduce dimensionality. Components contributing to the behavior were analysed separately as well.

### Manganese-Enhanced MRI analysis

#### Osmotic pump implantation

Osmotic mini-pumps (100 µL, Alzet) were filled with a solution of manganese(II) chloride tetrahydrate (MnCl₂·4H₂O, Sigma Aldrich) in the Tris buffer. Solution concentrations were individually prepared to achieve a dose of 60 mg/kg per animal. Pumps were primed overnight in a 37°C water bath prior to the surgical procedure. Vetalgin (200 mg/kg) was administered subcutaneously as an analgesic 30 minutes before surgery. Mice were anesthetized with isoflurane (2,5% induction, 1.5–2% maintenance) and secured in a stereotaxic frame. Meloxicam (0.5 mg/kg) was given immediately before the procedure, and Bepanthen eye ointment was applied to prevent corneal drying. Pumps were implanted subcutaneously on the dorsal surface beneath the scapula to avoid interference/restriction of animal movement. Postoperatively, all mice received Metacam in their drinking water for three days, followed by a seven-day monitoring with strict criteria for exclusion period prior to imaging (evaluation of surgery wound, body weight, food and water intake, behavior, mobility, and overall appearance).

#### Animal handling

Sedation was initiated using 2.5% isoflurane. Animals were fixed stereotactically in a prone position on an MR-compatible animal bed and maintained under inhalation anesthesia (isoflurane 1.5–2.5% in 100% oxygen, 1.5 L/min). To prevent drying of the eyes, Bepanthen eye ointment was applied. Body temperature was assessed using a rectal thermometer and maintained in the range of 36-38 °C throughout the experiment by means of a warm water heated silicon pad. Respiration was measured using a pressure sensor placed under the thorax. Depth of isoflurane anesthesia was adjusted to reach a respiration rate of 80 – 120 breaths per minute.

#### MR acquisition

MEMRI images were collected at 9.4 Tesla (BRUKER Biospec 94/20) in a cross-coil mode with radiofrequency transmission using a volume resonator, and room temperature 2×2 surface array coils for signal detection.

For localization, we ran a 2D FLASH sequence (TR = 525 ms, TE = 3.2 ms, flip angle 30°, 384 x 384 voxels, in-plane FOV 20 x 20 mm, 30 slices, 0.5 mm slice thickness, 0 mm slice gap). The scan time for this 2D FLASH was 2 min 31 sec. Then, a series of three 3D T1-weighted FLASH images was collected with TR = 20.1 ms, TE = 5.87 ms, flip angle 30°, matrix size 200 × 150 × 100 voxels, FOV 20 x 15 x 10 mm^3^, bandwidth 25 kHz, 2 averages, RF spoiling, read orientation along the rostral-caudal axis. Each of these three 3D images was acquired in 20 min 50 sec. Total acquisition time, including adjustments and localizer scans, was about 75 minutes.

BRUKER data were converted using the Python Bru2nii function. During this step, the nominal voxel size was enlarged by a factor of 10 to better match the SPM12 default values for human-sized brains in the following analyses.

#### Image preprocessing

Preprocessing steps were performed using MATLAB (with custom scripts), SPM12 (www.fil.ion.ucl.ac.uk/spm), 3D Slicer (vers. 5.6.2, www.slicer.org), and ANTs (Tustison et al., 2021).

The three individual runs per subject were realigned to the first image, and a mean image per subject was calculated. Mean images were co-registered to Hikishima template (28), and its tissue probability maps were utilized to segment all subjects (ANTs Atropos). Two (of the three) segments were used to produce individual brain masks that were subsequently refined by applying a slightly dilated mean group mask (to eliminate occasional imperfections of non-brain coverage). Rigidly aligned whole images were then first bias corrected using bias fields that were derived from the FLASH sequence, as these latter images showed fewer structural details of different tissue compartments (Slicer’s N4ITK routine) (29). Thereafter, all individual bias-corrected MEMRI images were brain-extracted (by applying previously produced individual brain masks) and warped to this group template space using ANTs registration. Finally, normalized images were spatially smoothed with a Gaussian kernel of 3 mm.

### cFOS immunohistochemistry and quantification

#### Brain extraction

Mice were subjected to CSDS for 10 days and were analysed for non-responders and responders using the social avoidance test (n=15 CSDS, 5 control). On the following day, mice were sacrificed 90min-120min after one round of defeat and brains were harvested using transcardial perfusion using 4% paraformaldehyde (PFA). Brains were then fixed overnight in 4% PFA and then in 30% sucrose in phosphate buffered saline (PBS) for 48 hours. Brains were stored in PBS at 4 °C till sectioning and immunohistochemistry.

#### Tissue sectioning and immunohistochemistry

Brains were selected at random from designated non-responders and responder groups for subsequent analysis. Brains (4 controls, 5 non-responders, 4 responders) were cryo sectioned using OCT (optimal cutting temperature compound) serially at 40micron using a sliding cryostat (Histo-line, MC 4000). Sections were stored in antifreeze solution (sucrose 30%, ethylene glycol 15%, Na-azide 0.02%, PBS). Immunohistochemistry was performed on free floating sections (1 in 4 whole brain series). First, sections were washed 3 times in PBS at RT (10 min each), and subsequently incubated in blocking solution at room temperature (1% PBS, Triton 0.3% and bovine serum albumin, 1%) for 90 minutes under constant shaking. Sections were then incubated in primary antibody solution against cFOS for two days at 4°C under constant shaking (1:5000, Synaptic System # 226 008 in PBS 1% and Tryton 0.1%). After a 20 min incubation at room temperature, sections were extensively washed in PBST and incubated with an Alexa conjugated secondary antibodies: Donkey anti-rabbit 647 (1:1000, Invitrogen, A31573) diluted in PBST 0.1 % at RT for 2 hours at room temperature under constant shaking. Sections were then washed 3 times with PBS, incubated with DAPI (1:1000 in PBS) for 5 min and mounted on superfrost glass slides (ThermoScientific) with Fluoromount (Invitrogen) after 2 additional washes in PBS. Images were acquired with a Zeiss Axioscan microscope Z1 equipped with a Hamamatsu Orca Flash 4 camera (2048×2048 pixels, 6.5 µm pixel size) with a 20x/0.8 Plan Apochromat objective. The resulting images’ pixel size is 0.325 µm, with 16-bit color depth.

#### Whole brain cFOS immunostaining image analysis

Whole-brain datasets were first registered to the 10µm Allen Brain Atlas (30) using ABBA (31). Slice positioning on the anterior-posterior axis and angle correction were manually performed. For all sections, Atlas alignments were first performed automatically concatenating affine and spline registrations (32,33) and subsequently manually refined using BigWarp (34) to maximize precision. For this, white matter landmarks and DAPI densities were used. Atlas registration accuracy was checked by at least two independent experimenters. All atlas annotations were imported into QuPath v0.5.1 (35). Quantification of cFOS+ cells was performed on 16-bit grey scale images using BraiAn (31) extension for QuPath. Positive cell detection, segmentation, and classification were performed following the procedures described in Chiaruttini et al. 2025. Damaged or misaligned tissue portions were excluded from further analysis.

### cFOS distribution analysis

#### cFOS density plot and module detection

cFOS signal densities for each brain structure and each mouse was analysed using an R script and method that has been described in detail in another publication (36). Data was distributed across brain subdivisions to a depth of up to 5 of the Allen Brain Atlas common framework. Bootstrapping (iteratively resampling with replacement) was performed for each group comparison and each brain structure with 10,000 repetitions in the R statistical environment. At the end, after accounting for missing values, a total of 142 brain regions were analyzed for cFOS distribution across the whole brains and all the groups. Volcano plot was utilized to visualize changes in brain regions among the groups by plotting fold change to the t-test p-value. Signal density in each of the brain regions was presented in the groups based on brain location using Pearson correlation. The brain modularity was assessed based on co-activation (presence of cFOS signal) with the aid of correlation matrix in the same region at the time of data sampling. Modules were identified based on the Euclidean distances between brain regions, calculated using the hierarchical clustering algorithm.

#### Network analysis

An analysis and graphical representation of the functional connections in the brain based on cFOS signal density correlation and modularity has been carried out based on a procedure detailed in a previous publication (36). To accommodate the large number of connections on one graph and narrow down the view to the most important relationships the network analysis was carried out using a positive correlation coefficient of value above 0.75. For the figure clarity, the connections in the whole brain network were shown using a threshold of 0.95.

The role of each of the regions in the network represented as nodes was calculated by module degree z-score (WMD, *connectivity inside local group*), participation coefficient (PC, *connectivity outside of local group*), and degree (k, *number of links with other nodes*).

### Chemogenetics based circuit manipulation and histology

To target the DR-BLA circuitry, chemogenetics/DREADD based inhibition was used. C57Bl/6n mice were injected with a combination of either an anterograde AAV containing inhibitory DREADD (AAV8-hSyn-DIO-hM4D(Gi)-mCherry) in the Dorsal Raphe with retrograde AAV (AAVrg.hSyn.HI.eGFP-Cre.WPRE.SV40) in the BLA or with anterograde control AAV (AAV5-Ef1a-DIO-mCherry) in DR with same retrograde AAV in the BLA, respectively (ADDGENE). The injection volume was 500nl of 1*10^12 ifu per region using stereotaxic surgery based on coordinates from Mouse Brain in Stereotaxic (DR −0.45, 0.6, 3.2; BLA −1.9, 3.3, 4.8). After surgery, mice were allowed to recover for 4-5 weeks following which they were subjected to the CSDS paradigm or housed as controls. After 10 days of CSDS, mice were given an intraperitoneal injection of Clozapine N oxide (CNO) at a concentration of 3mg/kg 30 min before behavioral assessment through Social Avoidance (day 11) or Elevated Plus Maze (day 12). Following the behavioral tests, mice were sacrificed and brains were harvested through intracardial 4% paraformaldehyde (PFA) perfusion for post hoc validation of surgery.

### Statistical Analysis

For SA and EPM, statistical analysis was carried out with Graphpad Prism using t-Test or one-way ANOVA with multiple comparison for single factor and two-way ANOVA with post hoc tests for mean comparison for two factor analysis. For DeepOF behavioral analyses, principal component analysis (PCA) was performed and R version 4.3.3 was utilized for assessing statistical significance for individual behavior component analysis. P<0.05 was selected as the significance threshold.

For MEMRI, statistical analysis was performed using a full factorial model in SPM12 with the three groups: controls, non-responders and responders. The global signal defined as summed intensities of all voxels in the brain was added as a regressor in order to compensate for possible interindividual differences in overall Mn^2+^ accumulation. We evaluated the effect of the CSDS intervention (i.e., control vs. non-responders and responders) as well as the stress susceptibility (i.e., non-responders vs responders) at a statistical threshold of p_uncorrected_<0.005, with a minimum cluster extent of 20 voxels. Partial least square analysis was conducted with the Braian python package for cFOS mapping. An R statistical package was used for cFOS data analysis details of which have been depicted previously in another publication (36).

## Results

### Deep phenotyping of behavioral trajectories and stress predisposition

To investigate the behavioral consequences of chronic stress, we employed a 10-day CSDS protocol, categorizing the mice into responders and non-responders, alongside unstressed controls. Mice were further profiled before and after the stress paradigm in a free social interaction paradigm using DeepOF. The experimental timeline is illustrated in Fig. 1A.

**Figure 1.**
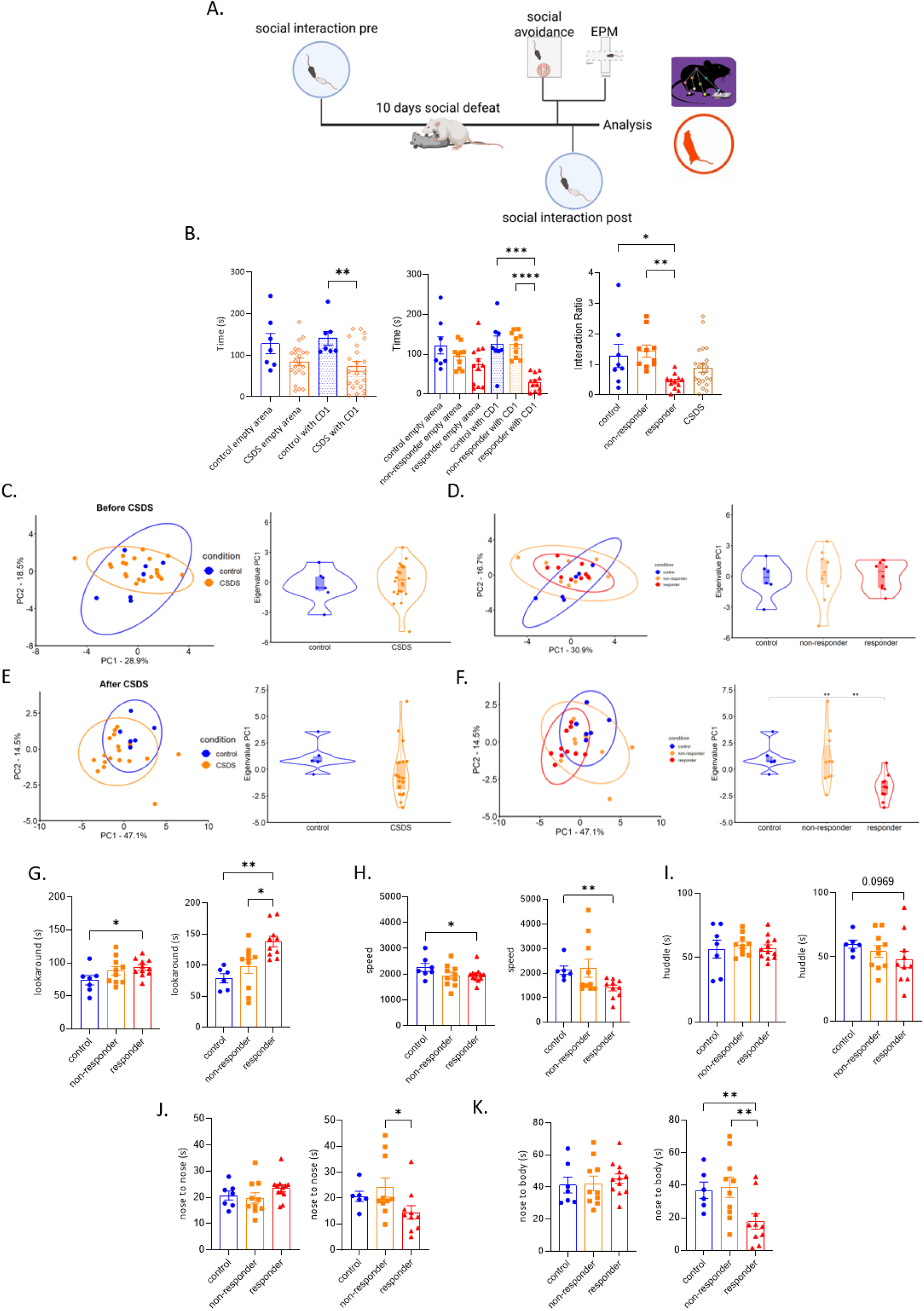
Classification criteria along with deep learning based pre and post stress (CSDS) behavioral analysis. **A.**Schematic of the experimental pipeline. Social Interaction test was conducted and recorded for deep learning based automated free social behavioral (DEEPOF) analysis before the starting of chronic social defeat protocol (CSDS) for 10 days. Elevated Plus Maze and Social Avoidance tests were conducted at the end of the CSDS protocol. A further round of Social Interaction test was carried out after this to be analysed by DEEPOF. **B.** Social Avoidance test was employed to classify responders and non-responders. Left panel: mice that underwent CSDS spent significantly less time in the interaction zone with CD1 mice present as compared to the controls and as compared to empty arena (without CD1) (Two way ANOVA within groups F(3,33)=5.623, p<0.01). Middle panel: responders spent significantly less time in the interaction zone with CD1 present as compared to controls and non-responders (two way ANOVA within groups F(5,54)=8.365; multiple comparison test p<0.01 and p<0.001). Right panel: Interaction ratio (time spent with CD1 vs without in the interaction zone) was significantly lower in responders as compared controls and non-responders (one way ANOVA F(3,48)=4.796, p<0.01; multiple comparison t-test p<0.05 and p<0.01). **C**. Free social behavioral (DEEPOF) analysis before CSDS. Left panel: PCA analysis shows distribution of mice before CSDS as compared to controls. Right panel: violin plot of PC1 eigenvalues pre-CSDS showed no significant differences between control and CSDS groups. **D.** DEEPOF analysis before CSDS. Left panel: PCA plot at baseline shows distribution of controls, non-responders and responders (based on classification post CSDS). Right panel: violin plot of PC1 eigenvalues showed no significant differences among the three groups – controls, non-responders and responders (based on classification post CSDS). **E.** DEEPOF analysis after CSDS. Left panel: PCA plot shows distribution of controls and mice that underwent CSDS. Right panel: violin plot of PC1 eigenvalues for mice that underwent CSDS vs controls showed no significant differences. **F.** Left panel: PCA plot for classified groups after CSDS. Right panel: Violin plot of PC1 eigenvalues revealed that responders were significantly different to controls and non-responders. **G-K**. Analysis of specific behaviors contributing to PC1 eigenvalue differences. Left panels are for before CSDS and right panels depict after CSDS for the same group of mice. **G.** Mice that were classified as responders showed significant differences in “lookaround” as compared to controls at baseline (one way ANOVA F(2,25)=2.777; multiple comparison t-test p<0.05). Post CSDS responders showed significant differences as compared to both controls and non-responders (one way ANOVA F(2,23)=8.985, p<0.05; multiple comparison t-test, p<0.01 and p<0.05). **H.** “Speed” of responders were significantly lower than controls pre-CSDS (one way ANOVA F(2,26)= 1.869, p>0.05 and multiple comparison t-test p<0.05), which became more prominent after the CSDS paradigm (one way ANOVA F(2,23) = 3.002, p=0.06 and multiple comparison, p<0.01). **I.** Time spent to “huddle” was not significantly different among the groups post CSDS though responders showed a trend of reduced huddle as compared to controls (one way ANOVA F(2,23)=1.078, p>0.05 and multiple comparison t-test p=0.0969). Similarly, no significant differences were observed among the groups before CSDS (one way ANOVA F(2,26)=0.19, p>0.05). **J.** There was a significant difference in “nose to nose” contact time between responders and non-responders after CSDS (one way ANOVA F(2,23) = 2.779, p= 0.08 and multiple comparison t-test, p<0.05) but not before CSDS (one way ANOVA F(2,26) = 1.383, p>0.05). **K.** Time spent by responders in “nose to body” was significantly lower as compared to the controls and non-responders after the CSDS (one way ANOVA F(2,23) = 4.716; p<0.05 and multiple comparison t-test, p<0.05). No significant differences were observed before CSDS (one way ANOVA F(2,26)=0.2754, p>0.05). * corresponds to p<0.05, ** corresponds to p<0.01, *** corresponds to p<0.005

As expected, the 10-day CSDS protocol elicited divergent behavioral responses, allowing for the classification of responders and non-responders based on Social Avoidance test (Fig. 1B). Notably, these groups showed no significant differences in the elevated plus maze (Fig. S1A), suggesting that the observed phenotypes were specific to the social challenge context rather than a generalized anxiety-like state. In the free social interaction test prior to CSDS, principal component analysis (PCA) of all detected behaviors revealed no significant differences between the later control and CSDS groups (Fig. 1C), nor between the mice subsequently classified as responders or non-responders (Fig. 1D). Following the CSDS paradigm, while no significant differences were observed in the PCA comparing control and CSDS animals (Fig. 1E), an analysis of the subgroups revealed that the responders were significantly different from both non-responders and controls (Fig. 1F).

To further characterize these differences, we assessed the individual behavioral categories which were most prominent between the groups before and after stress (Fig. S1B). Among these, ‘lookaround’ - a vigilance like behavior – was significantly altered in responders compared to non-responders and controls after CSDS. Interestingly the responder group exhibited significant differences in ‘lookaround’ behavior even at baseline, before the stress paradigm (Fig. 1G). A similar pattern was observed for ‘speed’, where responders differed significantly from controls both before and after stress exposure (Fig. 1H). In contrast, other behaviors as ‘huddle’ (Fig. 1I), which can be interpreted as an anxiety-like state, and ‘nose to nose’ (Fig. 1J) or ‘nose to body’ (Fig. 1K), which measure aspects of social interaction, only significantly differed after CSDS between responders vs non-responders and controls. These findings indicate that responders may possess a behavioral predisposition that is present at baseline and subsequently accentuated by stress.

### Mapping divergent whole brain activity patterns via longitudinal MEMRI

Behavioral changes in response to stress are often underpinned by systemic changes in brain activity. We hypothesized that responders and non-responders would exhibit distinct, brain-wide activity patterns. To capture these changes, we utilized Manganese-Enhanced MRI (MEMRI), taking advantage of the principle of manganese accumulation in high calcium flux cells (37). Following 10 days CSDS and subsequent classification into responders and non-responders, mice were implanted with subdermal osmotic pumps for continuous delivery of manganese chloride. After overnight rest, mice were subjected to a supervised modified social defeat paradigm (mCSDS) for the next 7 days, where the integrity of the implants was maintained, before MRI imaging. A schematic of the experimental pipeline is depicted in Fig. 2A.

**Figure 2.**
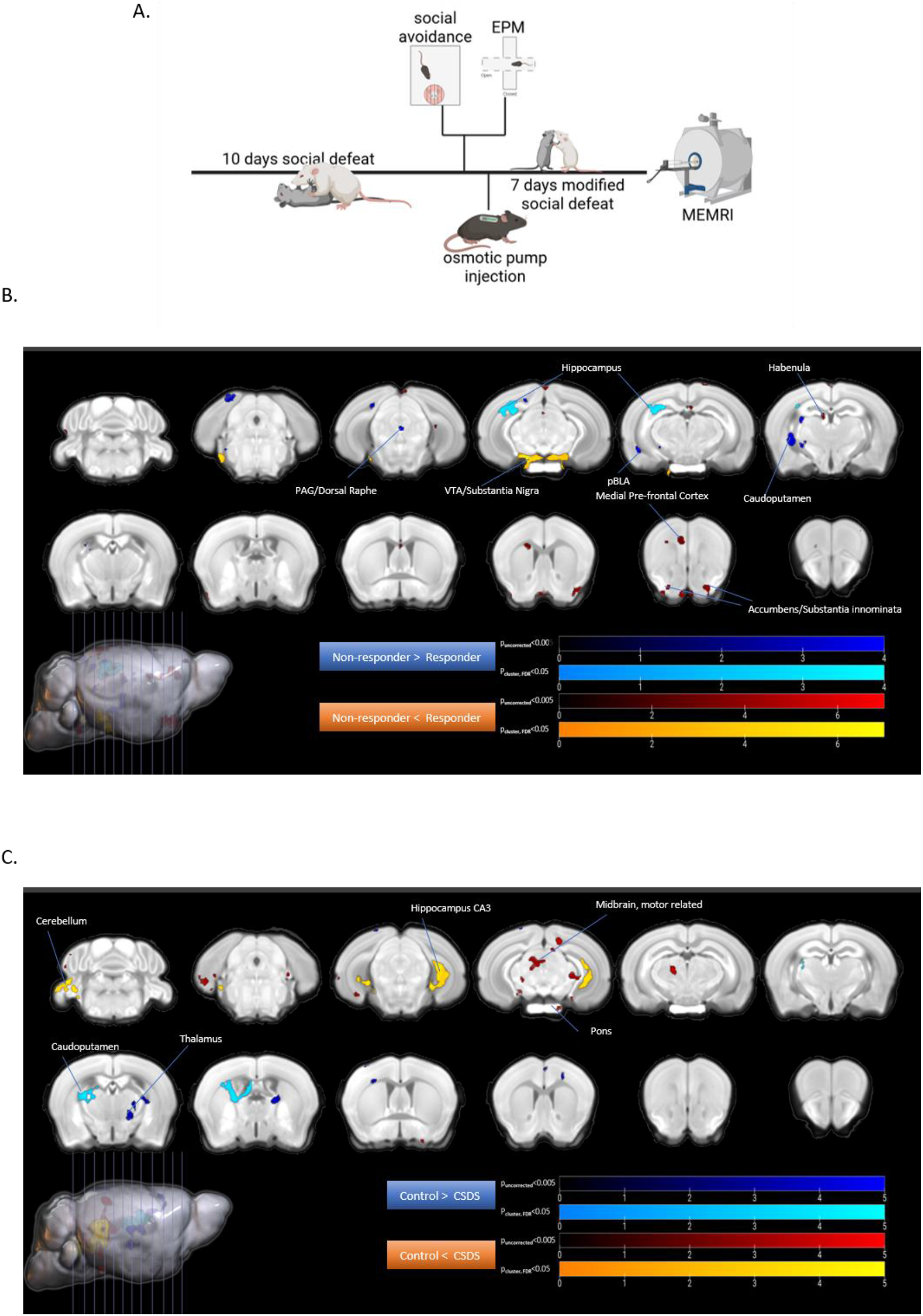
Manganese (Mn)-Enhanced MRI to elucidate differences between non stress controls, non-respondents and respondents. **A.** Schematic of the experimental pipeline. Post 10 days social defeat (CSDS), mice were classified as responders and non-responders based on Social Avoidance tests. Osmotic pump carrying Manganese Chloride (MnCl2) solution was placed sub-dermally. This was followed by one day of recovery and 7 days of modified social defeat and subsequently MRI analysis. **B.** Plates showing differences in multiple regions where the Mn intensity, and thus, activity was significantly different in the responders as compared to the non-responders. **C.** Plates showing differences in different regions where the Mn intensity, and thus, activity was significantly different in the CSDS group (comprising of responders and non-responders) as compared to the controls. Statistical T-maps (threshold p<0.005) have been superimposed onto mouse brain Atlas (Allen). n=8 controls, 7 non-responders and 8 responders

MEMRI analysis revealed multiple regions that showed differential activity between responders and non-responders (Fig. 2B), as well as between the unsegregated CSDS group and controls (Fig. 2C). Specifically, the divergence between responders and non-responders was characterized by differential activation in the periaqueductal grey (PAG), DR, hippocampus (HIP), ventral tegmental area/substantia nigra (VTA/SN), medial prefrontal cortex (mPFC), habenula and nucleus accumbens. In contrast, the comparison between CSDS animals and controls identified differentially activated regions including, cerebellum, hippocampus, thalamus and midbrain.

### Whole-brain cFOS mapping reveals functional network architecture

Next, to investigate how chronic stress reconfigures global brain plasticity, we performed brain-wide cFOS mapping following a social challenge. This approach enabled us to identify differentially affected regions and characterize the emergent functional networks associated with each behavioral strategy. After the CSDS paradigm and subsequent identification of responder and non-responders, mice were subjected to a final social defeat and sacrificed two hours later for global cFOS analysis (31). A schematic of the experimental protocol is shown in Fig. 3A, with representative cFOS stainings in Fig. S2.

**Figure 3.**
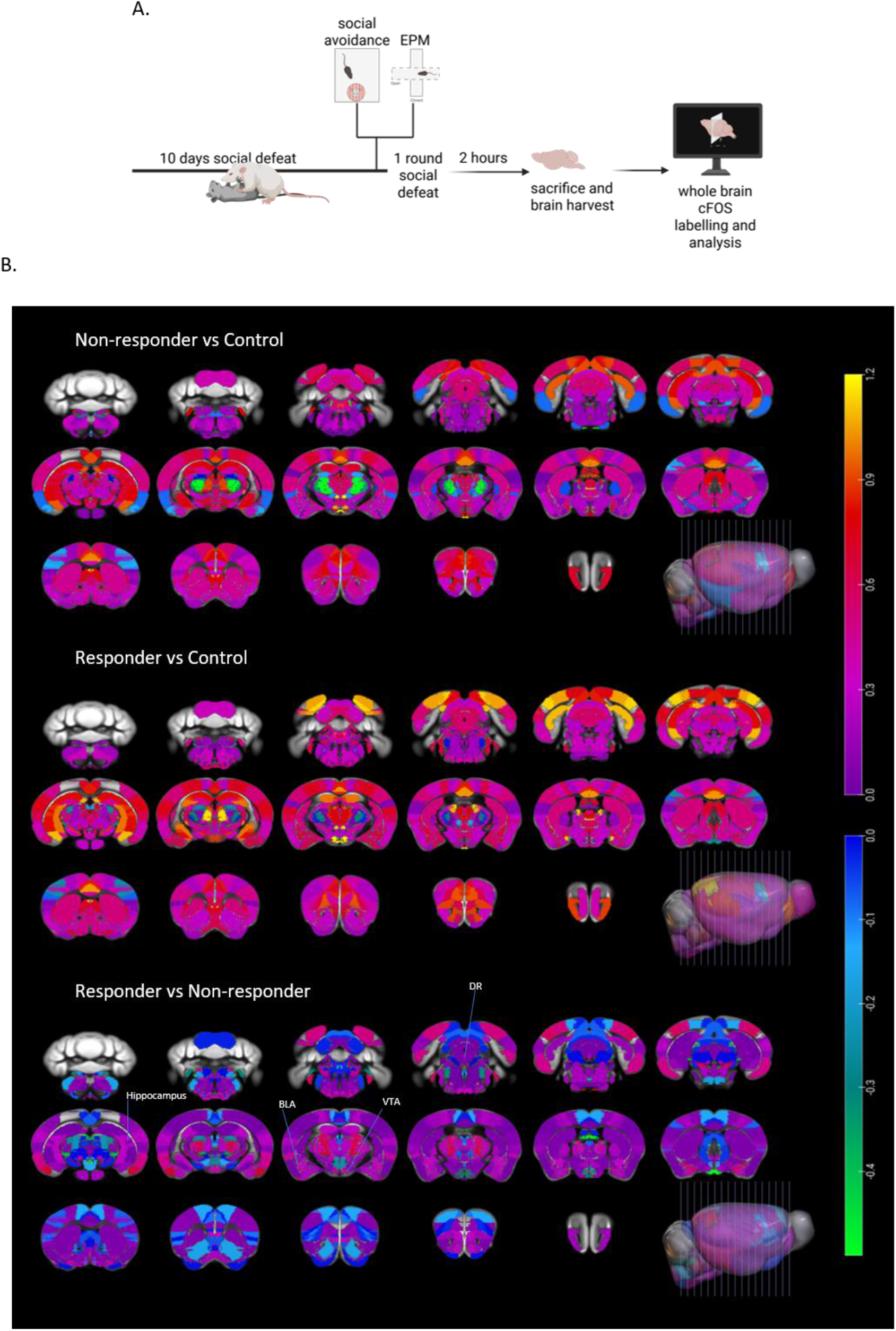
Whole brain c-FOS mapping to find differentially activated regions amongst non-stressed controls, non-responders and responders. **A.** Schematic of the experimental procedure. Mice underwent 10 days of social defeat (CSDS) followed by classification into responders and non-responders based on Social Avoidance tests. After one further round of social defeat, mice were sacrificed after 2 hours and brains were harvested for subsequent analysis using anti cFOS labelling. **B.** Brain map showing differences in cFOS density in different brain regions compared across the groups. n=4 controls, 4 non-responders and 5 responders.

Regional density analysis revealed a distinct divergence in neural activation patterns. A global heatmap of cFOS densities across brain regions (Fig. 3B; Table S1A) and results distribution on volcano plot (Fig. 4A) demonstrated that responders exhibited a significantly higher number of regions with >1.5-fold changes compared to controls than did non-responders.

**Figure 4.**
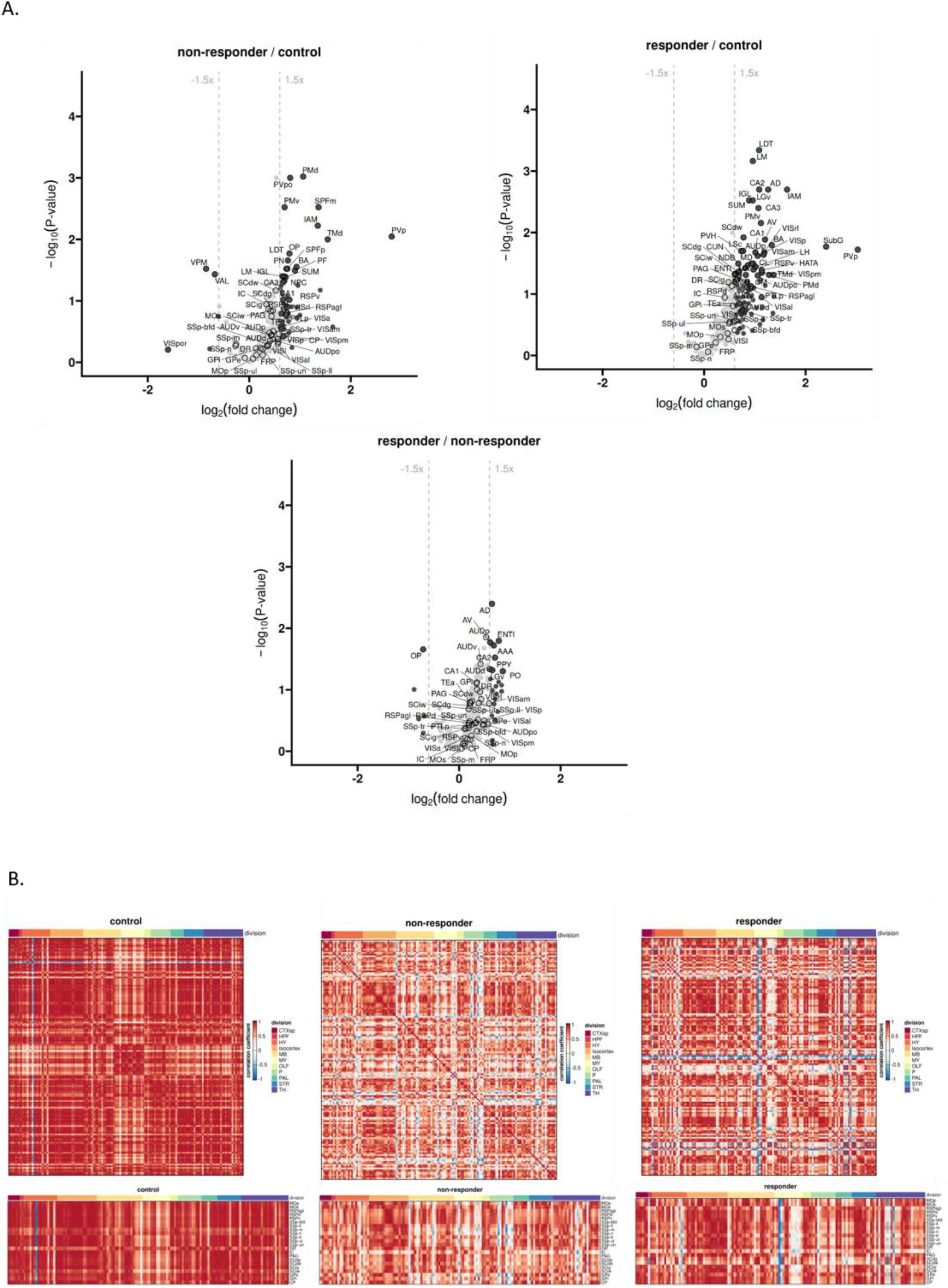
Analysis of distribution of cFOS across different brain regions after CSDS. **A**. Volcano plots of T test of P values (significance) and fold changes (sqrt) of regions showing differences in non-responders vs control (top left), responders vs controls (top right) and responders vs non-responders (bottom). **B**. Correlation heatmaps of different brain regions categorised into major divisions according to Allen Brain Atlas for responders (top left), non-responders (top right) and controls (bottom). R values represented as cool to warm colors depicts increased correlation of signal densities between two regions, respectively. In each figure a correlation plot of all the regions is accompanied by a plot of cFOS densities of cortical areas and few selected areas as dorsal raphe (DR), peri-aquaductal grey (PAG) across all divisions of brain analysed.

To understand how these regional changes translate into system-wide alterations, we performed correlation and network analysis (Fig. 4B, Tables S2-5) comparing cFOS densities in different brain regions distributed across major brain divisions (Fig. 4B upper panel). Inter-brain region correlations were clearly reduced in animals with a history of chronic stress compared to controls, independent of their coping strategy. Interestingly, responders displayed higher cFOS correlations particularly in the isocortex, midbrain, olfactory lobes and parietal lobes compared to non-responders. A highly similar pattern emerges when the analysis is directed to brain regions that were identified to be differentially activated in the MEMRI analysis, including the DR (Fig. 4B lower panel).

To identify the specific nodes driving these global network shifts and compare with existing knowledge, we narrowed our analysis of selected regions that have previously been reported to contribute to stress system regulation and that have been observed in our MEMRI experiments (Fig. 5A). These plots clearly depicted that cFOS densities in the BLA, baso-medial amygdala (BMA), CA2 of the hippocampus, DR and paraventricular nucleus of the hypothalamus (PVH) of the responders were significantly higher than non-responders and controls, whereas CA1 and CA3 of the hippocampus showed significant increase in cFOS immunoreactivity in both responder and non-responder groups as compared to controls. These findings were further validated using bootstrapping to ensure the robustness of the regional differences (Fig. S3, Table S16).

**Figure 5.**
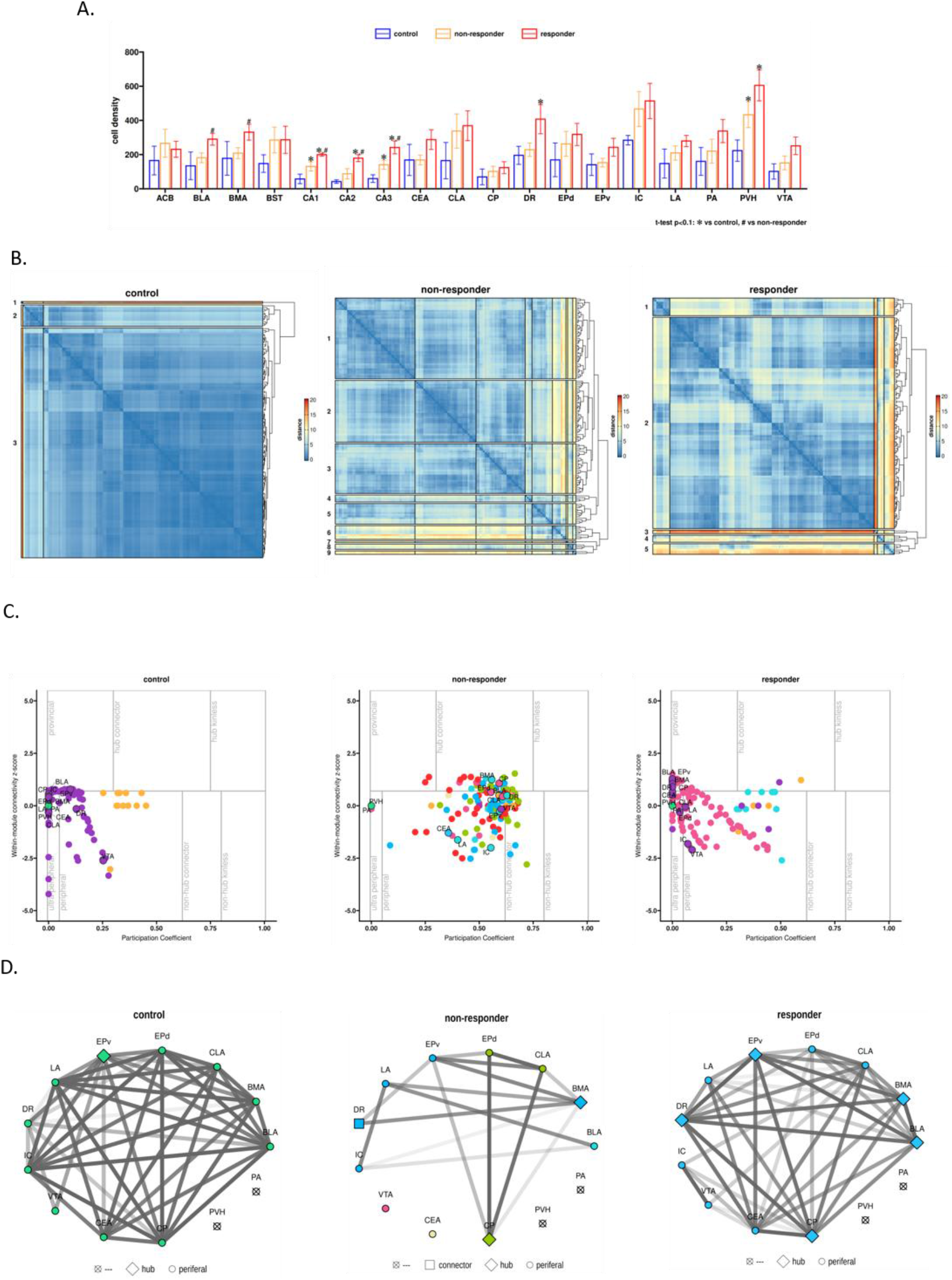
Network analysis of plasticity changes measured by cFOS density. **A**. Bar plots for cFOS cell densities in selected brain regions among the three groups. **B**. Module analysis of closely associated regions based on co-expression and correlation of cFOS densities in responders, non-responders and controls. Depicted in each of the matrices are relative distances between brain regions in a hierarchical organisation within the clustering of sub-networks in each groups. They are arranged in ascending order in accordance to their distances in sub-dendogram, with shorter Euclidean distance depicted by warmer color. Increased number of modules indicate higher heterogeneity and complexity of associated regions. **C**. Type of nodal connections within module connectivity - plots are based on parameters for each region in the modular network to the z score of the modular degree. **D**. Network map among selected regions of the brain in responders, non-responders and controls showing differential connectivity. Thicker edges indicate higher correlation. A threshold of 0.95 has been used for the plots. Each of the nodes have been further classified as connector, hub or peripheral based on strength of association. Connectors indicate groups with higher connectivity outside local group, whereas hubs indicate higher local connectivity.

Next, we applied modular analysis to understand the changes in architecture of the networks (Fig. 5B) (36,38–41). This clustering analysis revealed a striking difference in network architecture, with a higher modularity in the non-responder group (total 9) as compared to the responders (total 5) and controls (total 3). In controls, nodes remained closely associated with low Euclidean distances, reflecting a highly integrated, unchallenged system. The higher modularity in experimental groups, especially non-responders suggested greater functional segregation, a configuration often associated with system resilience to localized perturbations but at the cost of fluid information flow between modules (Fig. 5B). Consistent with this, non-responders showed a higher number of hub-connectors—nodes that facilitate inter-module communication—whereas responders were characterized by a higher proportion of provincial nodes (Fig. 5C).

Interestingly in the responders, key stress-related regions including PAG, BLA, DR, and VTA were all clustered inside a single module (Module 4). In contrast, these regions were segregated into distinct modules in non-responders (e.g., BLA and VTA in Module 5; PAG, DR, and IC in Module 9). This clustering in responders suggests that the coordinated activity of these regions may underpin the stress-responsive phenotype, whereas their segregation in non-responders may reflect a more rigid, resistant network state.

We next employed analysis to examine the functional connectivity of brain regions (Fig. 5C and 5D, Tables S6-15). This resulted in classification of the regions as connectors, hubs or peripheral nodes, depending on connections within or out of the module they belong (Fig. S4). The analysis of brain region roles in the network, revealed that control group consisted mostly of peripheral nodes, without clear central function in the communities. In the responder group, brain regions were mostly peripheral or provincial with more within-module connections, and few hub connectors with more links within and out of their module. The most striking change were noted in non-responders, where many regions had relatively higher participation coefficient linking numerous modules in the network, and higher within module degree.

A functional network connection map representing selected regions identified as differentially activated in the cFOS or MEMRI analysis (Fig. 5D) revealed that for many of these regions the intensity of functional network connectivity was weaker in the non-responders compared to responders and controls. In particular a functional network connection of DR to BLA was lost specifically in the group of non-responder animals.

All these analyses have also been conducted without segregating the groups into responders and non-responders and can be found in Fig S5A-F. As expected, we observed a significantly higher number of regions exhibiting greater than 1.5-fold changes in cFOS density as compared to the control along with reduced correlation among the regions (Fig. S5A, S5B). The modular analysis (Fig. S5C, S5D) revealed the existence of 10 distinct modules in contrast to controls, non-responders and responders (Fig. S5E). In the network analysis (Fig. S5F), we found a weaker but existing functional network between DR and BLA. While this consolidated analysis showed the impact of CSDS stress, it also indicated the importance of segregation of the group into responders and non-responders to prevent masking of data pertaining to individual variability while simultaneously exhibiting the strength and confidence of our analytical approach.

### Chemogenetic inhibition of the DR-BLA pathway promotes stress unresponsiveness

Our multi-modal analysis provided a convergent map of the neural substrates underlying stress responsiveness, highlighting the DR to BLA circuit as potential key node. While these regions have been implicated in aversive and valence behavior (42,43), their coordinated interaction as a discrete functional circuit in the context of individual stress adaptation was so far not described. Based on the high cFOS density of these regions and strong functional coupling specifically in the responder group, we hypothesized that the DR-BLA pathway serves as a critical driver of a rigid stress response phenotype.

To test this hypothesis, we employed a pathway-specific chemogenetic approach to selectively inhibit DR neurons projecting to the BLA. We injected a retrograde AAV-Cre vector into the BLA, while the DR received a Cre-dependent inhibitory DREADD or a control virus (Fig. 6A). Following recovery period, mice were subjected to 10 days CSDS and tested for social and anxiety behavior following acute CNO-mediated silencing of the DR-BLA circuit. Post-hoc immunohistological analysis confirmed successful expression of the viral constructs (Fig. 6B). Acute chemogenetic inhibition of the DR-BLA pathway significantly attenuated the social avoidance characteristic of the responder phenotype (Fig. 6C). Specifically, stressed mice expressing the inhibitory DREADD spent significantly more time exploring the interaction zone in the presence of the CD1 social target compared to stressed controls. Furthermore, circuit inhibition exerted a potent anxiolytic effect in the EPM, with DREADD-expressing stressed mice spending significantly more time in the open arms compared to stressed controls (Fig. 6D). These findings demonstrate that the DR-BLA pathway is a primary mediator of stress-induced adaptive behavior and inhibition of this circuit promotes a stress non-responsive phenotype.

**Figure 6.**
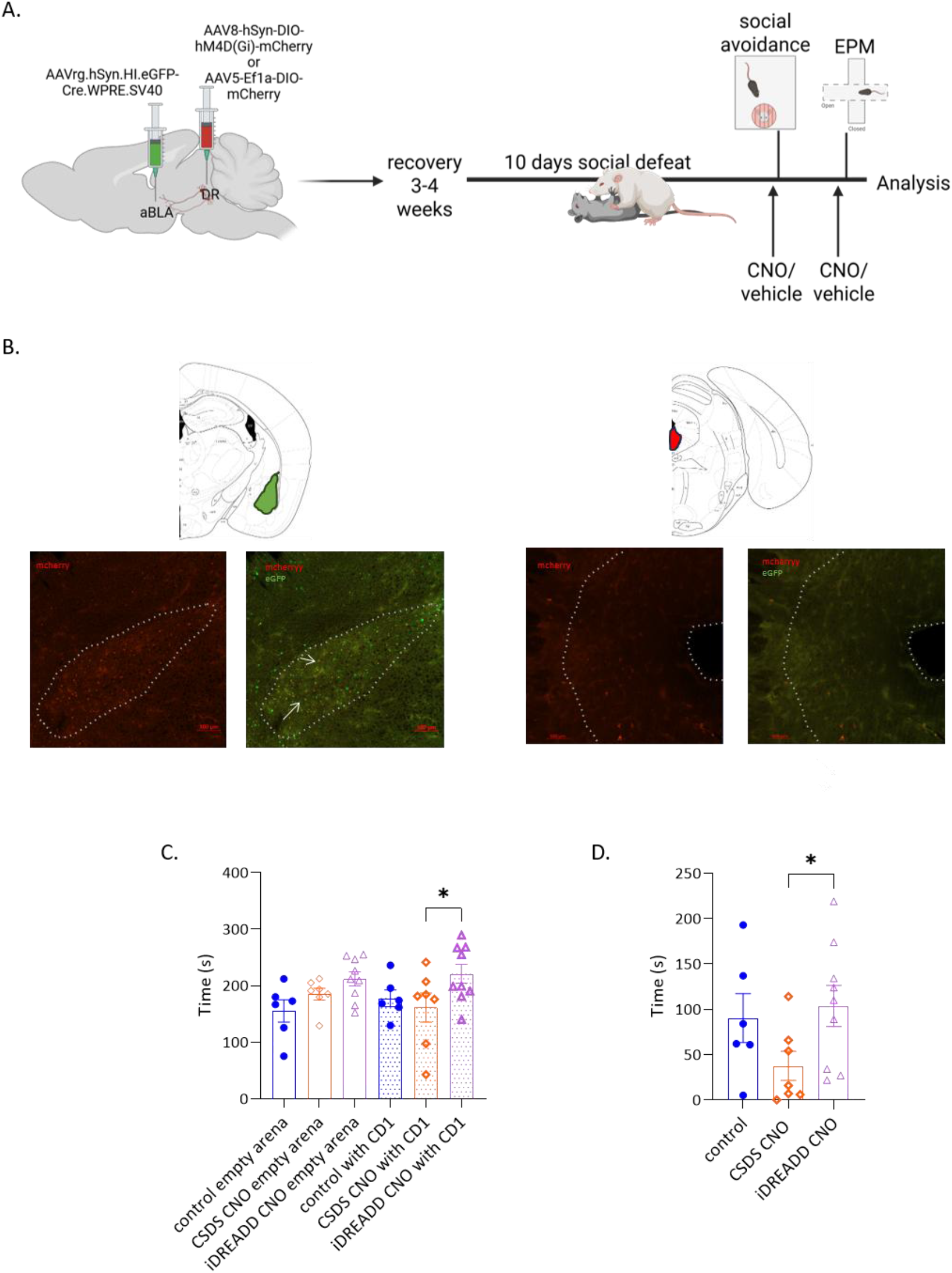
Chemogenetics based inhibition of DR-BLA (dorsal raphe – basolateral amygdala) circuitry to investigate its importance in responders vs non-responders. **A.** Schematic for the experimental procedure. Mice were injected with a retrograde virus AAVrg.hSyn.HI-eGFP-Cre.WPRE.SV40 in the anterior BLA and an anterograde virus (either an inhibitory DREADD carrying virus AAV8-hSyn.DIO-hm4D(Gi)-mCherry or a control AAV5-EF1a-DIO-mCherry) –thus labelling neurons specifically in the DR-BLA circuitry. After a recovery phase of 3-4 weeks, mice underwent 10 days of social defeat (CSDS) protocol. They were tested on the Elevated Plus Maze and Social Avoidance tests on consecutive days with an injection of Clozapine N oxide (CNO) or vehicle (Saline) 30 min prior to experimental procedure. **B.** Depiction of regions shown reference to Mouse Brain in Stereotaxic co-ordinates. (59) Representative pictures depicting expression of the virus in BLA (left panel) and in DR (right panel). Scale 100um **C.** Social Avoidance test-Mice expressing inhibitory DREADD upon CNO injection, spent significantly more time in the interaction zone with the CD1 mice present as compared to stressed (CSDS) groups and controls (two-way ANOVA within groups F(5,30)=3.429, p<0.05; multiple comparison t-test p<0.05). **D.** Elevated Plus Maze test-Mice expressing inhibitory DREADD upon CNO injection spent significantly more time in the open arms of the maze as compared to the CSDS group (One way ANOVA F=2.490, p=0.1095; multiple comparison t-test p<0.05). n= 6 (controls), 7 (CSDS) and 9 (iDREADD). * corresponds to p<0.05, ** corresponds to p<0.01, *** corresponds to p<0.005

## Discussion

The current study demonstrates that divergent stress-coping strategies are driven by distinct, system-wide reconfigurations of brain activity and functional plasticity. By pairing machine-learning-based deep behavioral phenotyping with longitudinal MEMRI and whole-brain cFOS mapping, we identified divergent behavioral coping trajectories following chronic social stress, which correspond to distinct brain network activity changes. Crucially, our network-level approach identified a conserved DR-BLA circuit as a primary orchestrator of these phenotypes.

A central question of our study is the re-evaluation of how individual variability is classified following chronic stress. Traditional rodent stress frameworks frequently interpret animals that exhibit social avoidance as “susceptible” or “vulnerable,” while framing non-avoidant animals as “resilient” (7). However, displaying avoidance toward a previously encountered, highly aggressive conspecific can also be seen as a fundamentally normal, adaptive, and expected learned response to threat (44). Here, rather than viewing this behavior through the lens of pathology, our approach distinguishes between actively adapting individuals (“responders”) and those displaying a rigid, non-responsive phenotype (“non-responders”). To capture this difference beyond simple proximity metrics, we leveraged machine-learning-based deep phenotyping via DeepOF (23). Similar to our previous observations using a prolonged CSDS paradigm, stress elicited a number of significant behavioral differences in free social interaction domains, indicating behavioral adaptations as heightened vigilance and reduced social contact. Strikingly, mice that ultimately transitioned into the active responder phenotype displayed distinct behavioral kinetics, like specifically accelerated locomotive speed and heightened vigilance-like “lookaround” behavior, at baseline, prior to any stress exposure. These pre-stress behavioral signatures suggest a latent neurobehavioral predisposition, reinforcing previous findings that pre-existing traits or endophenotypes dictate subsequent stress reactivity (45–47). Ultimately, high-dimensional behavioral analysis demonstrates that active stress adaptation is a complex, partially pre-configured behavioral trajectory, rather than a simple deficit in social approach.

To map the functional topography underlying these phenotypes, we utilized Manganese-enhanced MRI during chronic stress exposure. This approach revealed a distinct divergence between responders and non-responders across multiple brain regions, including the PAG/DR, basolateral amygdala (BLA), hippocampus (HIP), ventral tegmental area (VTA/SN), lateral habenula (LH), medial prefrontal cortex (mPFC), and nucleus accumbens/substantia innominata. These structures form the core of canonical valence and stress-regulatory circuits (7,48). Crucially, MEMRI captures consolidated, enduring network activity reconfigurations while filtering out transient, acute stress responses. Consequently, we detected robust differences predominantly when segregating chronically stressed animals in responders and non-responders, as the unsegregated analysis masks individual-specific neural signatures. Despite variations in experimental design and imaging modalities, our data converge with previously published results identifying regions as the hippocampus, caudoputamen, hypothalamus, periaqueductal grey and cerebellum as differentially activated between unsegregated CSDS animals and controls (49). More importantly, the phenotypic divergence we observed in the ventral hippocampus, VTA/SN, mPFC and nucleus accumbens aligns with recent resting-state fMRI functional connectivity networks associated with divergent stress coping strategies (17). This consistency across distinct imaging techniques and stress paradigms reinforces the biological validity of our findings and confirms these macro-scale networks as robust, conserved substrates of individual stress trajectories of responders and non-responders.

While our MEMRI framework captured the accumulated, chronic imprint of stress over time, cFOS profiling allowed us to interrogate the dynamic responsiveness and immediate activation of the brain-wide neural circuitry under acute stress. Network and modularity analyses of these expression patterns revealed profound system-level reconfigurations unique to each behavioral strategy. Specifically, non-responders exhibited marked hyper-modularity (nine modules) compared to responders (five modules) and unstressed controls (three modules). This structural fragmentation indicated a high degree of functional segregation, a configuration that limits information flow across the global network and likely underpins the rigid, non-responsive behavioral phenotype. While global cFOS analysis exhibited multiple regions that have been previously associated with variability in stress response like the BLA, HIP, VTA and hypothalamus (7,11,50), our network approach revealed how these nodes dynamically couple or desynchronize. For example, our analysis identified the claustrum (CLA) as an area that demonstrated differential functional connectivity changes in responders compared to non-responders, in line with previous reports highlighting this region’s role in stress-induced emotional processes (51). Strikingly, core emotional and valence-processing nodes, including the PAG, VTA, BLA, and DR, co-clustered into a single, tightly integrated functional module in responders, but were segregated into isolated modules in non-responders. Our network connectivity analysis presented several interesting novel altered functional couplings among different brain regions. Most notably, among others, functional coupling between the DR and the BLA was entirely lost in non-responsive animals. This stark divergence demonstrated that active stress adaptation requires highly coordinated, inter-modular communication, pointing directly to the DR-BLA axis as one critical, unexamined gatekeeper that may be dictating individual behavioral trajectories.

To directly establish causality, we functionally interrogated the DR-BLA circuit using a pathway-specific chemogenetic approach. While the DR is a classical modulator of stress coping primarily via its serotonergic neuronal population (52–54), and the DR-BLA circuit has been implicated in broader memory processes (43,55), its necessity in driving individual behavioral coping strategies under chronic stress conditions remained unexplored. Acute silencing of BLA-projecting DR neurons during a social challenge attenuated social avoidance and reversed anxiety-like behavior. By functionally decoupling this circuit, we successfully shifted the active behavioral adaptation of responders toward a rigid, non-responsive phenotype, demonstrating that the strong inter-modular functional connectivity observed in our cFOS data is a requirement for active stress coping. A key consideration for future mechanistic dissection is the profound cellular heterogeneity that characterizes both the DR and the BLA (56–58). Disentangling the contributing neuronal populations will be essential to map exactly how distinct cellular sub-circuits drive either active adaptation or rigid unresponsiveness under chronic stress.

In summary, our findings offer a powerful blueprint for decoding individual stress trajectories within a framework centred on active adaptation versus behavioral rigidity. By pairing machine-learning-based deep phenotyping with brain-wide activity and plasticity mapping, we demonstrate that the divergence between responders and non-responders reflects distinct, system-wide network reconfigurations. Crucially, our causal validation of the DR-BLA circuit as a modulator of these phenotypes highlights a specific circuit mechanism dictating stress coping strategies. Moving forward, leveraging this framework to resolve precise cellular heterogeneities and sex-specific circuits will be essential to define the neurobiology of individual stress-coping differences, ultimately driving more precise, personalized therapeutic strategies for stress-related conditions.

## Acknowledgements

The work has been carried out at the Max Planck Institute of Psychiatry and was funded by Max Planck society and, by DFG grant JU 3039/5-1(BJ), and NCN grant 2023/51/B/NZ4/01135 (MB, MP, MS). We thank Ms Daniela Harbich at MPIP for her assistance in experimentation. We thank the Animal House Facility at Max Planck Institute of Psychiatry for their help in animal maintenance and experimentation. We express our gratitude to Ms Michaela Blažíková of the Institute of Molecular Genetics of the Czech Academy of Sciences and EuroBioImaging for aiding in carrying out cFOS analysis. The laboratory of BAS is supported by the European Research Council (ERC-2021-STG 101042309), the Fondazione Cariplo (2020-3632), Airalzh (AGYR2021), the Alzheimer’s Association (AARG-22-974392) and the Université Côte d’Azur. The graphics in the manuscript have been designed in BioRender and MS Powerpoint.

## Data Availability

The whole brain cFOS analysis data is available upon request.

## Competing Interests

The authors declare no conflicts of interest.

## Authors Contributions

SM and MVS conceptualised, designed the study and wrote the manuscript. SM carried out experiments, performed interpretation and analysis and co-ordinated among contributors. MVS gave critical inputs in interpretation and refined the analysis. TS and MC aided in MEMRI experimentation and imaged the mice. TS performed analysis on MEMRI data. BAS, BDB, CC and MP contributed in global cFOS labelling, imaging and analysis. MB and MS carried out global cFOS analysis. SN, CB and SM carried out DeepOF analysis. REH and SM carried out surgical interventions. MP, LvD, JB, MS, YH, VK, LA, BJ, AR contributed directly in carrying out the experiments. NS supported in acquiring microscopy data.

## Notes

### Competing Interest Statement

The authors have declared no competing interest.

