## Supplementary Figures for "Deep Phenotyping with Global Brain Activity and Plasticity Mapping Identify the Dorsal Raphe-Basolateral Amygdala Circuit as a Mediator of Adaptive Stress Responses"

Figure S1. *Behavioral analysis of non-stress control vs CSDS stressed and responders vs non responders*. **A**. Left panel - Elevated Plus Maze analysis of cohort used for DEEPOF analysis revealed no significant differences in time spent in open arms between controls and CSDS (t-test p>0.05). Right panel- no significant differences were also found comparing responders to non-responders and controls (one way ANOVA F(2,27)=1.389, p>0.05 and multiple comparison p>0.05) **B**. Top contributing behaviors in PCI for controls, non-responders and responders (in Fig 1d and f) before (left panel) and after (right panel) CSDS . **C,D.** Behavioral analysis of a second cohort of mice used in MEMRI experiment. **C.** Social Avoidance test. Left panel- mice that underwent social defeat (CSDS) exhibited a reduced time in the interaction zone with CD1 present as compared to the controls (ANOVA groups F(3, 36)= 3.321, p<0.05; multiple comparison t-test p<0.05). Middle panel- responders spent significantly lesser time in the interaction zone with CD1 mice present as compared to the controls and non-responders (ANOVA between groups F(5,34)=11.21, p<0.001; multiple comparison t-test p<0.01 and p<0.001). Right panel- the interaction ratio for the responders were also significantly lower than the non-responders (one way ANOVA F(3,32)=3.753, p<0.05; multiple comparisons p<0.05). **D.** Elevated Plus Maze. Left panel -there was no significant differences in time spent in the open arms by the CSDS group as compared to the controls (t-test p>0.05). Right panel - No significant differences were observed between the responders and the non-responders in terms of time spent in open arms (One way ANOVA F(2,17)=0.5588, p>0.05; multiple comparisons t-test p>0.05).

* corresponds to p<0.05, ** corresponds to p<0.01, *** corresponds to p<0.005, **** corresponds to p<0.001

Figure S2. Representative images of slides for whole brain c-FOS analysis showing c-FOS distribution in red punctas in controls, non-respondents and respondents

Figure S3. *Bootstrapping of cFOS density data from whole brain among all groups.* Bootstrapping estimates differences that deviate significantly from observed difference between groups indicating less precise results. Bootstrapping has been carried out with 10,000 repetitions by iteratively resampling observed data. A significant deviation was observed for the following structures – VPM, VAL, VPL, V, SF, SGN.

Figure S4. Network maps generated from correlation matrices among different brain regions in case of controls (top), non-responders (middle) and responders (bottom). Thicker edges indicate higher correlation. A threshold of 0.95 has been used for the plots. Each of the nodes have been further classified as connector, hub or peripheral based on strength of association. Connectors indicate groups with higher connectivity outside local group, whereas hubs indicate higher local connectivity.

Figure S5. Comparison of CSDS group to control for whole brain c-FOS mapping. **A**. Volcano plots of T test of P values (significance) and fold changes (sqrt) of regions showing differences in CSDS vs controls. **B**. Correlation heatmaps for brain regions for the CSDS (stressed) group among all the regions (top) and selected (below). **C**. Modules arising from clustering of sub-networks depicted in a matrix with heirarchial manner with relative Eucleadian distances in CSDS group (shorter distance is warmer color) **D**. Type of nodal connectivity within module connectivity in CSDS group **E**. Number of modules per group as depicted in Figure 5c and Suppl Fig 6d resulting in cutting the dendogram at 50% tree heights. **F**. Network connectivity maps among different regions for CSDS group (top) and selected (below). The nodes represent brain regions while the edges correspond to correlation coefficient. A threshold of 0.95 has been used for the images of these plots. Connectors indicate groups with higher connectivity outside local group, whereas hubs indicate higher local connectivity.

Figure S1

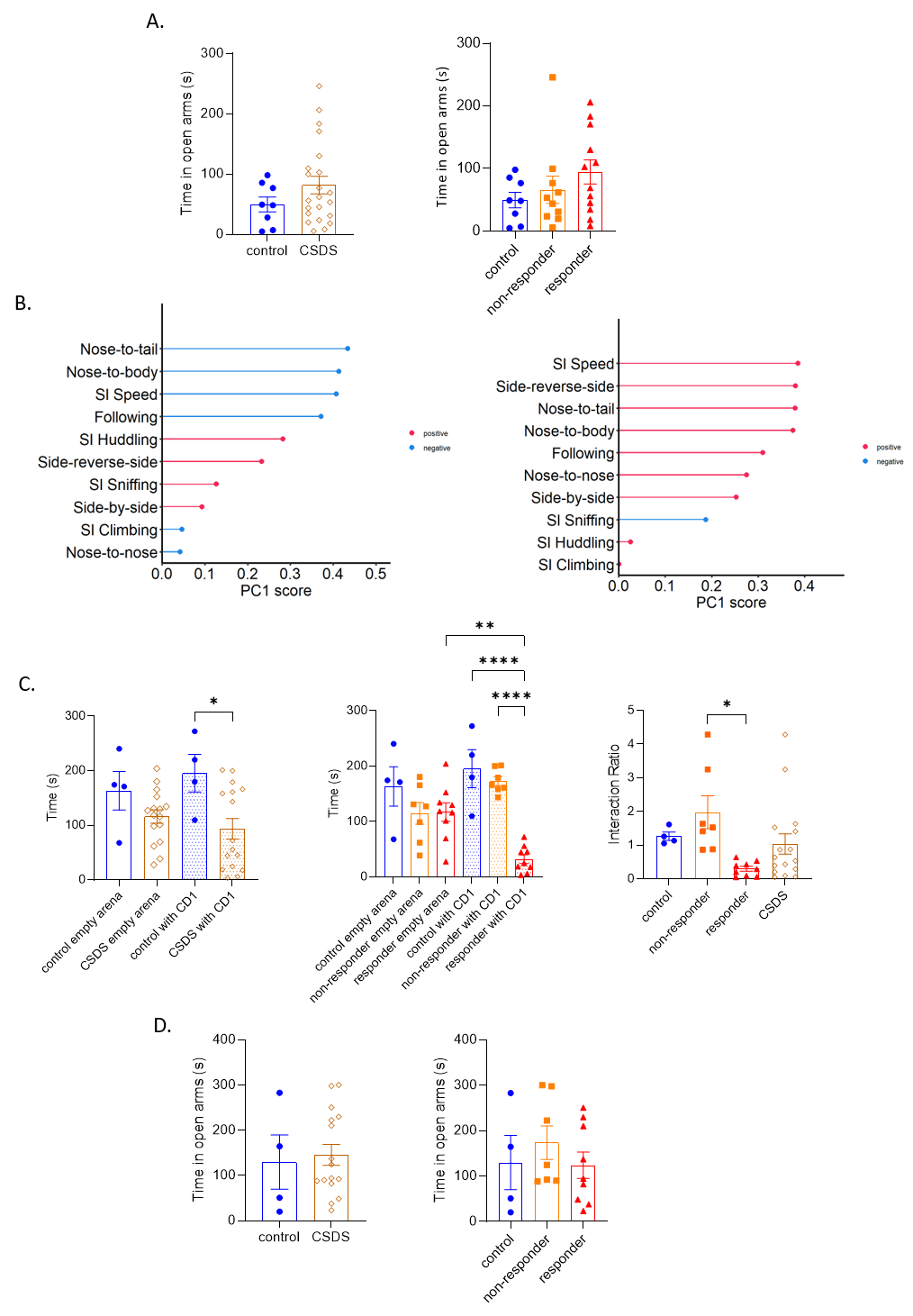

Figure S2

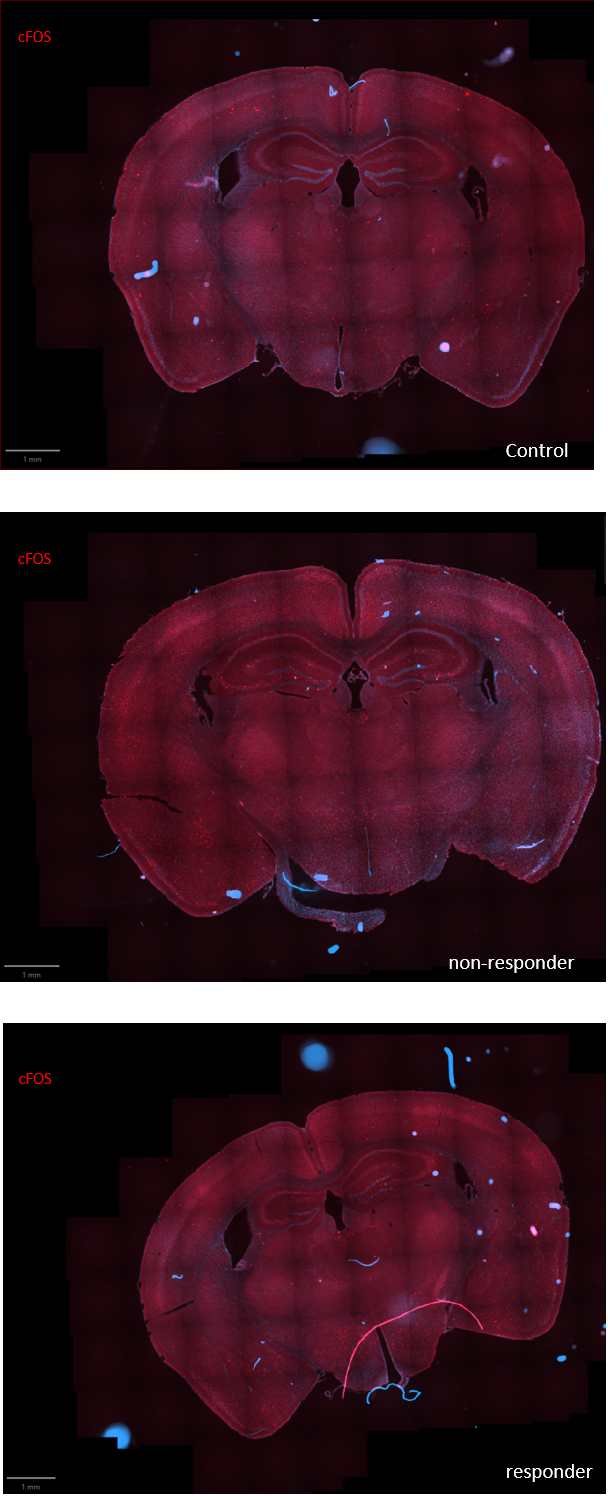

Figure S3

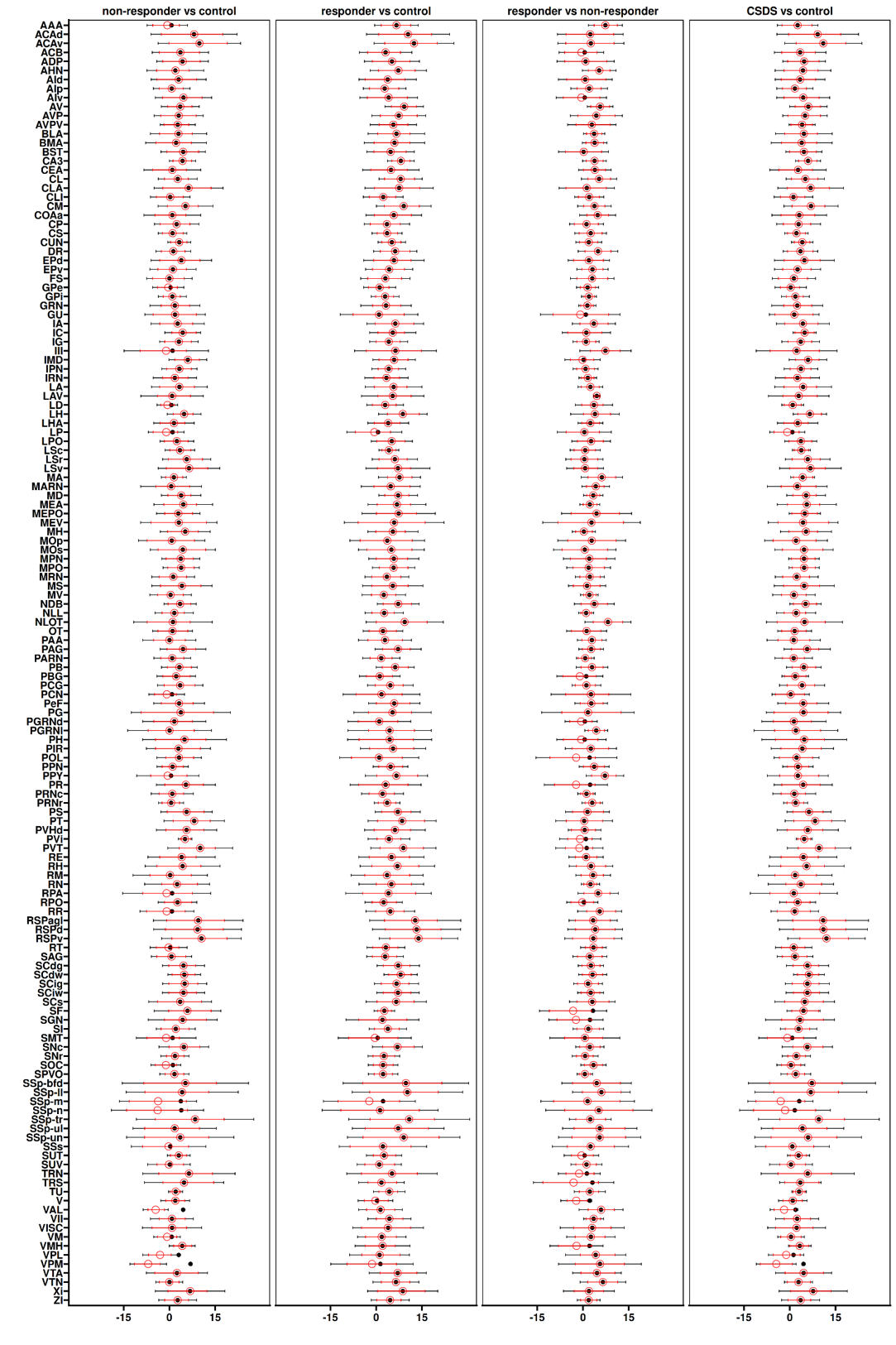

Figure S4

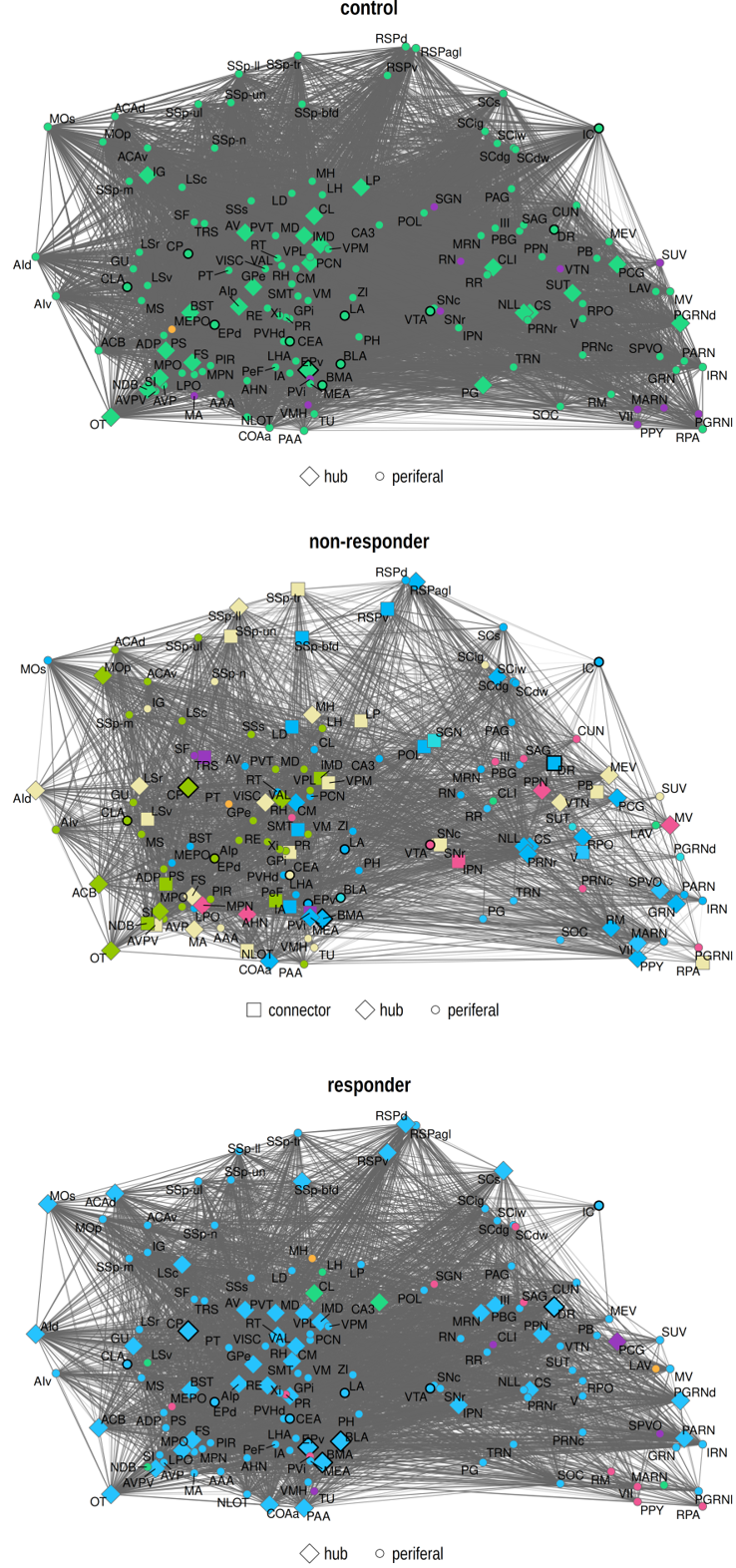

Figure S5

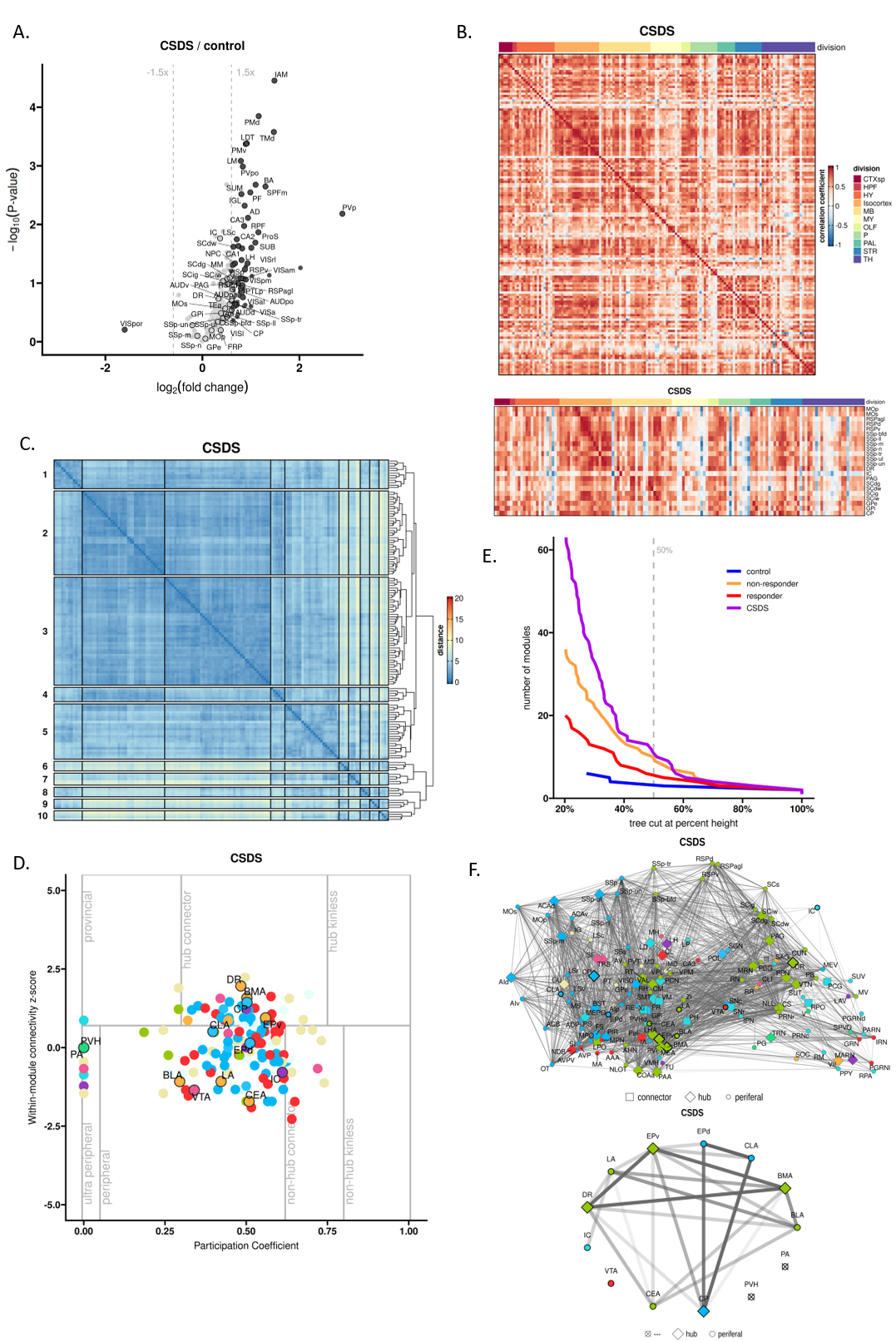
